# xSiGra: Explainable model for single-cell spatial data elucidation

**DOI:** 10.1101/2024.04.27.591458

**Authors:** Aishwarya Budhkar, Ziyang Tang, Xiang Liu, Xuhong Zhang, Jing Su, Qianqian Song

**Affiliations:** Luddy School of Informatics, Computing, and Engineering, Indiana University Bloomington; Department of Computer and Information Technology, Purdue University, Indiana, USA; Department of Biostatistics and Health Data Science, Indiana University School of Medicine, Indiana, USA; Gerontology and Geriatric Medicine, Wake Forest School of Medicine, North Carolina, USA; Department of Health Outcomes and Biomedical Informatics, College of Medicine, University of Florida, Florida, USA

**Keywords:** Explainable AI, spatial cell recognition, hybrid graph transformer, interpretable features

## Abstract

Recent advancements in spatial imaging technologies have revolutionized the acquisition of high-resolution multi-channel images, gene expressions, and spatial locations at the single-cell level. Our study introduces xSiGra, an interpretable graph-based AI model, designed to elucidate interpretable features of identified spatial cell types, by harnessing multi-modal features from spatial imaging technologies. By constructing a spatial cellular graph with immunohistology images and gene expression as node attributes, xSiGra employs hybrid graph transformer models to delineate spatial cell types. Additionally, xSiGra integrates a novel variant of Grad-CAM component to uncover interpretable features, including pivotal genes and cells for various cell types, thereby facilitating deeper biological insights from spatial data. Through rigorous benchmarking against existing methods, xSiGra demonstrates superior performance across diverse spatial imaging datasets. Application of xSiGra on a lung tumor slice unveils the importance score of cells, illustrating that cellular activity is not solely determined by itself but also impacted by neighboring cells. Moreover, leveraging the identified interpretable genes, xSiGra reveals endothelial cell subset interacting with tumor cells, indicating its heterogeneous underlying mechanisms within the complex cellular communications.

## INTRODUCTION

Recent advances in spatial transcriptomic techniques have enabled commercially available platforms for measuring mRNA expression in a tissue at molecular level spatial resolution, allowing biologists to gain novel insights about diseases^1^. Spatial locations of expressions are critical in understanding of cell-cell interactions and cell functioning in tissue microenvironment^2, 3^. Molecular imaging-based in-situ hybridization approaches such as NanoString CosMx Spatial Molecular Imaging (SMI), an automated microscope imaging system, allows spatial in-situ detection on formalin-fixed paraffin embedded samples. CosMx SMI is capable of detecting both RNAs and proteins on the same tissue slide, allowing 3-dimensional subcellular resolution image analysis with an accuracy of ∼50nm in the XY plane and high throughput (up to 1 million cells per sample)^4^. MERSCOPE^5^, another commercial platform, utilizes the MERFISH^6^ (multiplex error-robust fluorescence in situ hybridization) platform that captures hundreds to thousands of RNA specifies at the same time. Other noticeable commercial platforms that provide single-cell or sub-cellular spatial resolution include the 10x Genomics’ Xenium system and BGI’s SpaTial Enhanced REsolution Omics-Sequencing (Stereo-seq) in situ sequencing system using the DNA Nanoball technology.

The emerging spatial imaging technologies require tailored computational methods for data analysis, clustering, and enhancement. Some deep learning (DL)-based methods have been proposed for spatial transcriptomics data^7^. For example, the CCST^8^ method, designed based on Graph Convolutional Neural Network (GCN), takes both gene expression data and spatial information of single cells as input, to cluster spatial transcriptomics data. The SiGra^9^ method enhances the transcriptomics data and identifies spatial domains from the spatial multi-modality data. SpaGCN^10^ provides gene expression, spatial information, and histology information to the GCN network to identify spatial domains and spatially variable genes. stLearn^11^ is developed for spatial transcriptomics to identify cell types, reconstruct cell trajectories, and detect microenvironment. BayesSpace^12^ uses Bayesian approach and spatial information to enhance gene expression signals. STAGATE^13^ combines gene expression and cell locations to train a graph autoencoder for low-dimensional embedding and spatial domain detection. Other methods such as Seurat^14^ and Scanpy^15^ also provide functionalities for spatial transcriptomics data analysis and domain identification. Clustering methods such as Louvain^16^ perform domain recognition using expression data but do not utilize spatial information.

Although the above DL-based methods prove to identify spatial cells or domains with high accuracy, their intrinsic black-box nature inhibit the explainability, regarding what genes and cells are used by these methods to achieve accurate spatial identities^17^. Such explainability issue is common when applying advanced deep learning approaches^18^. Explaining the model decisions can aid to find any limitations and validate model functioning using known knowledge^19^. Several works are done to make the graph neural network models interpretable. For example, Saliency^20^ assigns the square of gradients as importance scores for the input features. Guided backpropagation^21^ uses only positive gradients for backpropagation while setting negative gradients to 0. Those positive gradients at the input layer are used as the importance score of input features. InputXGradient^22^ assigns importance score for input features as gradients multiplied by input. Deconvolution^23, 24^ computes the gradient of output class with respect to the input but propagates only the non-negative gradient and uses that as an importance score for input features.

In this study, we have proposed a novel method, i.e., xSiGra, to identify interpretable features contributing to the identification of spatial cell types. xSiGra takes advantage of the multi-modality spatial data including multi-channel histology images, spatial information, and gene expression data. xSiGra not only demonstrates its superior performance than existing state-of-the-art methods in the identification of spatial cell types, but also provides explainable information about the importance of cells and genes for the identification. The explainable capability of xSiGra also shows better performance than existing several graph-explainable algorithms, which facilitates biological insights from spatial imaging data.

## RESULTS

### Overview of the xSiGra model

The primary function of the xSiGra model is to elucidate the significance of each gene for biological annotations, such as cell identities, within high-resolution spatial data. As depicted in the overall diagram (**Fig.1**), cells of the same type exhibit diverse gene expression patterns and interact with their environment differently across various spatial locations and microenvironments. We illustrate this conceptually through two types of endothelial-stromal interactions: firstly, via growth factors secreted by stromal cells and receptors expressed on endothelial cell surfaces, and secondly, through the expression of specific structural proteins in the extracellular matrix (ECM) by stromal cells and the endothelial integrins responsible for recognizing and binding to these proteins.

**Fig 1:**
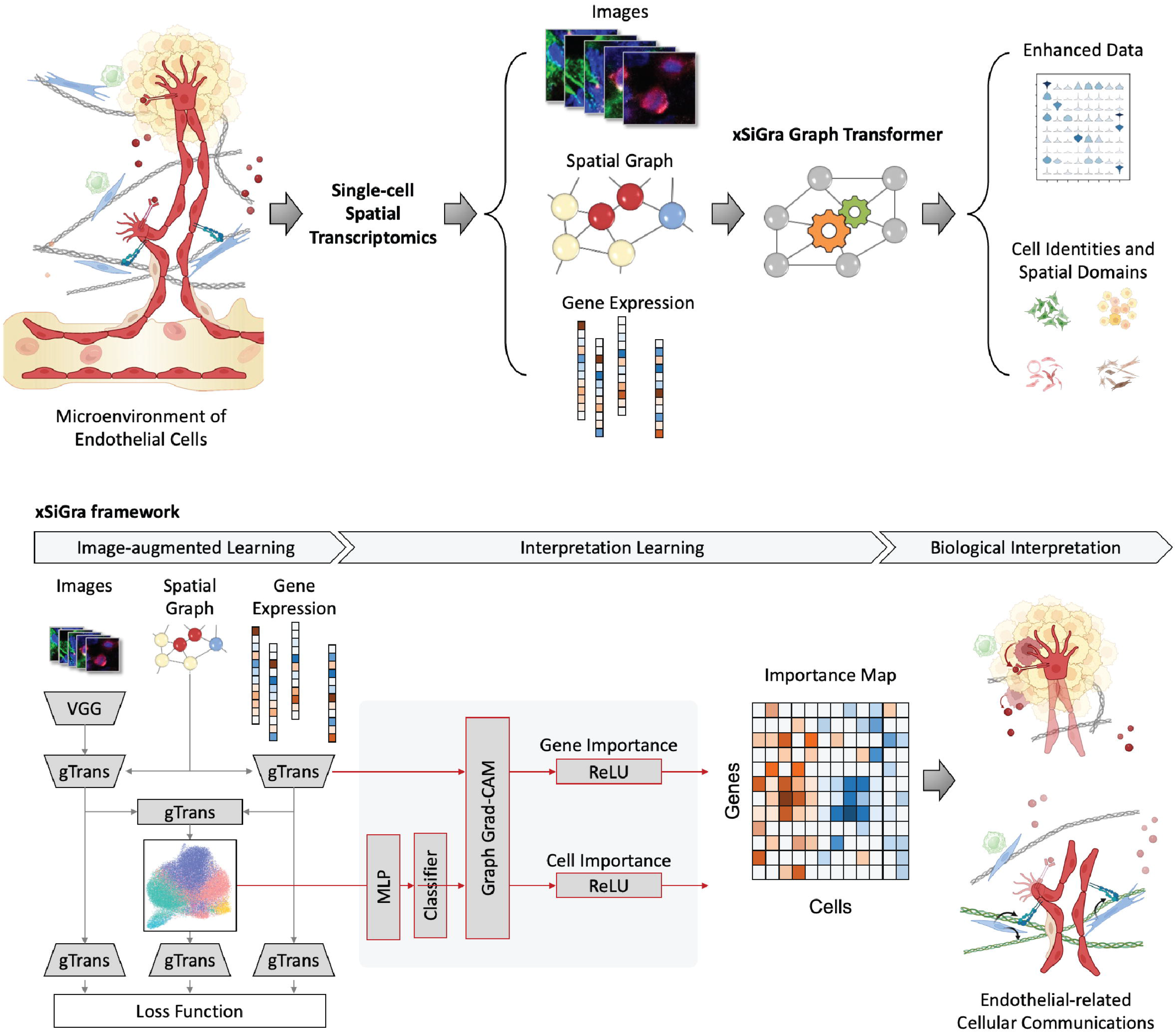
Overview of xSiGra model. **a** xSiGra uses spatial transcriptomics data to gain insights about cellular microenvironment. Spatial transcriptomics profiles are represented by graph structure with each cell in spatial graph having associated gene expression and multi-channel images. xSiGra generates enhanced gene expressions and identifies spatial cell types, with related interpretable features for biological insights. **b** xSiGra performs image augmented learning to enhance transcriptomics data (SiGra+) and recognize spatial cell types. It also provides interpretable features that can be used for biological interpretation.

The xSiGra model is composed of a SiGra+ module and an explainable module. The explainable module uses a novel graph gradient-weighted class activation mapping (graph Grad-CAM) algorithm to understand the contribution of each gene for different cell types within the spatial slides. The SiGra+ module learns the latent representation of multi-modal spatial transcriptomics data, which is used by the explainable module to explain which spatial gene expression patterns are used by SiGra+ for spatial cell type identification. The SiGra+ module also further enhances the original SiGra^9^ model by using a VGG feature extractor^25^ to improve histochemistry image feature extraction, and by introducing a Kullback–Leibler (KL) divergence loss^26^ to better balance the contributions from the transcriptomics and the imaging data.

### xSiGra accurately identifies spatial cell types in various datasets

We have compared our method with seven state-of-the-art methods, i.e., SiGra^9^, SpaGCN^10^, stLearn^11^, BayesSpace^12^, STAGATE^13^, Seurat^14^, and Scanpy^15^. Here we use 8 NanoString CosMx lung tissue slices (See **Data availability**) for performance comparisons. The adjusted rand index (ARI) score is used for evaluation of the detected spatial cell types, including tumor, fibroblasts, lymphocyte, myeloid, mast, neutrophil, endothelial, and epithelial cells. **Fig.2a** shows the ARI results of each of the methods across 8 lung cancer tissues. xSiGra achieved a median ARI of 0.565, better than SiGra (ARI = 0.505), stLearn (ARI = 0.360), Seurat (ARI = 0.315) and BayesSpace (ARI = 0.265), as well as STAGATE (ARI = 0.245), Scanpy (ARI = 0.235) and SpaGCN (ARI = 0.20). Specifically for Lung-13 tissue, which consists of 20 field of views (FOVs) and 77643 cells, **Fig.2b** presents the spatial cell types identified by different methods. The cell types identified by xSiGra match well with the ground truth with an overall ARI of 0.64, higher than SiGra (ARI = 0.45), stLearn (ARI = 0.60), Seurat (ARI = 0.35), Scanpy (ARI = 0.32), STAGATE (ARI = 0.29), BayesSpace (ARI = 0.28), and SpaGCN (ARI = 0.26). **Fig.2c** shows the spatial cell types at the FOV level for better clarity. The cells identified by xSiGra show consistency with the ground truth, identifying the continuous tumor region along with infiltrated immune cells. On the contrary, in FOV1, BayesSpace mislabels some lymphocytes and fibroblast as mast and endothelial cells while stLearn identifies some tumor cells and fibroblast incorrectly as mast and epithelial cells. In FOV5, BayesSpace misrecognizes some tumor cells as neutrophils; and fibroblasts are mislabeled as mast cells. stLearn mixes fibroblasts with neutrophil cells, as well as tumor cells with mast cells.

**Fig 2:**
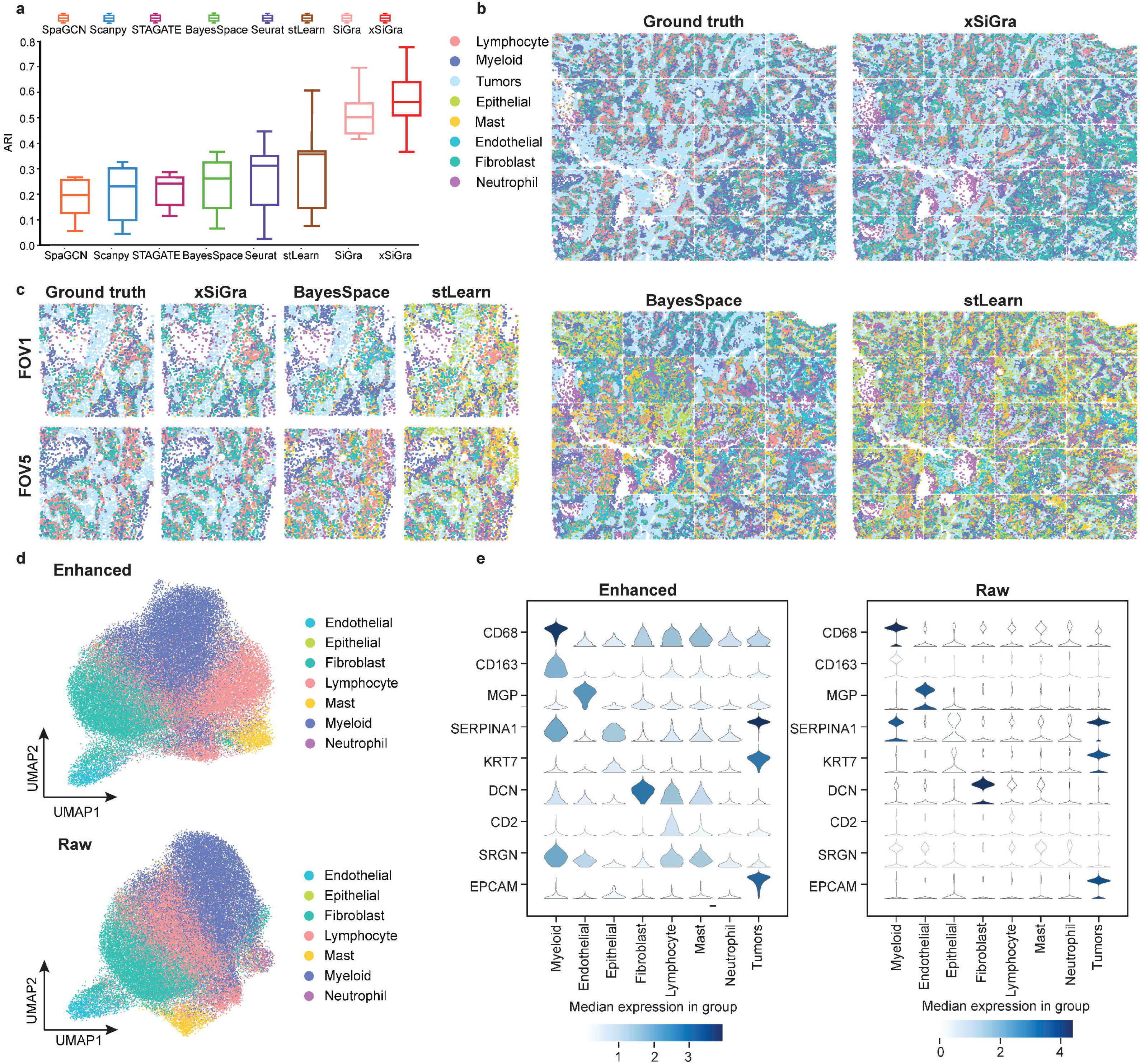
Performance evaluation on lung cancer slices. **a** Box plot of adjusted rand index for 8 NanoString SMI datasets is shown for each of the methods, i.e. SpaGCN, Scanpy, STAGATE, BayesSpace, Seurat, stLearn, SiGra, and xSiGra. The middle line represents the median and limits are first and third quartiles. **b** Spatial cell types of ground truth and those identified by stLearn and BayesSpace are shown. **c** Spatial cell types for 2 FOVs of ground truth and those identified by stLearn and BayesSpace are shown. **d** UMAP visualizations for raw and enhanced gene expressions are shown for non-tumor cells. **e** Violin plot of raw and enhanced gene expressions of marker genes in different cell types.

In addition to single-cell spatial data, xSiGra also proves to perform better in spot-based spatial data such as 10x Visium spatial slices. As shown in **Supplementary Fig.1a**, xSiGra achieves a higher ARI score compared to the state-of-the-art methods across 12 Visium datasets (See **Data Availability**). **Supplementary Fig.1b-d** present the spatial domains accurately identified by xSiGra. This demonstrates that in comparison with the existing methods, our method can more accurately identify the spatial domains in spot-based spatial data, and distinguish the different domains in the complex microenvironment. Moreover, xSiGra also enhances gene expression data for downstream analysis. From the Uniform Manifold Approximation and Projection (UMAP) analysis on enhanced data and raw data, enhanced data presents clearer separation of different cell types than raw data (**Fig.2d**). The enhanced gene expression data is further supported by the cell type markers whose expressions are highly expressed in their corresponding cell types (**Fig.2e**).

### xSiGra presents better interpretability for explainable features

Our method uses a novel graph Grad-CAM to identify the important cells and genes for spatial cell types. The model interpretability can be evaluated with two key metrics: fidelity and contrastivity^27^. Fidelity measures how much the detection accuracy is reduced when the important genes are excluded. Contrastivity measures the differences in important genes identified across different spatial cell types. Based on these two metrics, we have compared xSiGra with four existing explainable methods, including Saliency^20^, InputXGradient^22^, GuidedBackprop^21^, and Deconvolution^23^ on each of the lung tissue slices (See **Data Availability**). Specifically, **Fig.3a** shows the fidelity and contrastivity scores of each of the methods on the Lung-13 tissue slice, where xSiGra achieved a higher median fidelity score of 0.16, which is the best compared to Saliency (median: 0.10), InputXGradient (median: 0.09), GuidedBackprop (median: 0.05) and Deconvolution (median: 0.04). xSiGra also obtained better contrastivity score (median: 0.77) than the other methods, especially Deconvolution (median: 0.18). **Fig.3b** shows that the fidelity and contrastivity scores of different methods on this Lung-5 rep3 tissue slice. Across all eight lung cancer tissue slices (**Fig.3** and **Supplementary Fig.4**), xSiGra presents superior performance in both fidelity score (median = 0.16) and contrastivity score (median = 0.76), compared to the other methods (Saliency: 0.09, 0.45; InputXGradient: 0.08, 0.46; GuidedBackprop: 0.06, 0.16; and Deconvolution: 0.05, 0.23). Collectively, xSiGra demonstrates superior performance of interpretability than other explainable methods.

**Fig 3:**
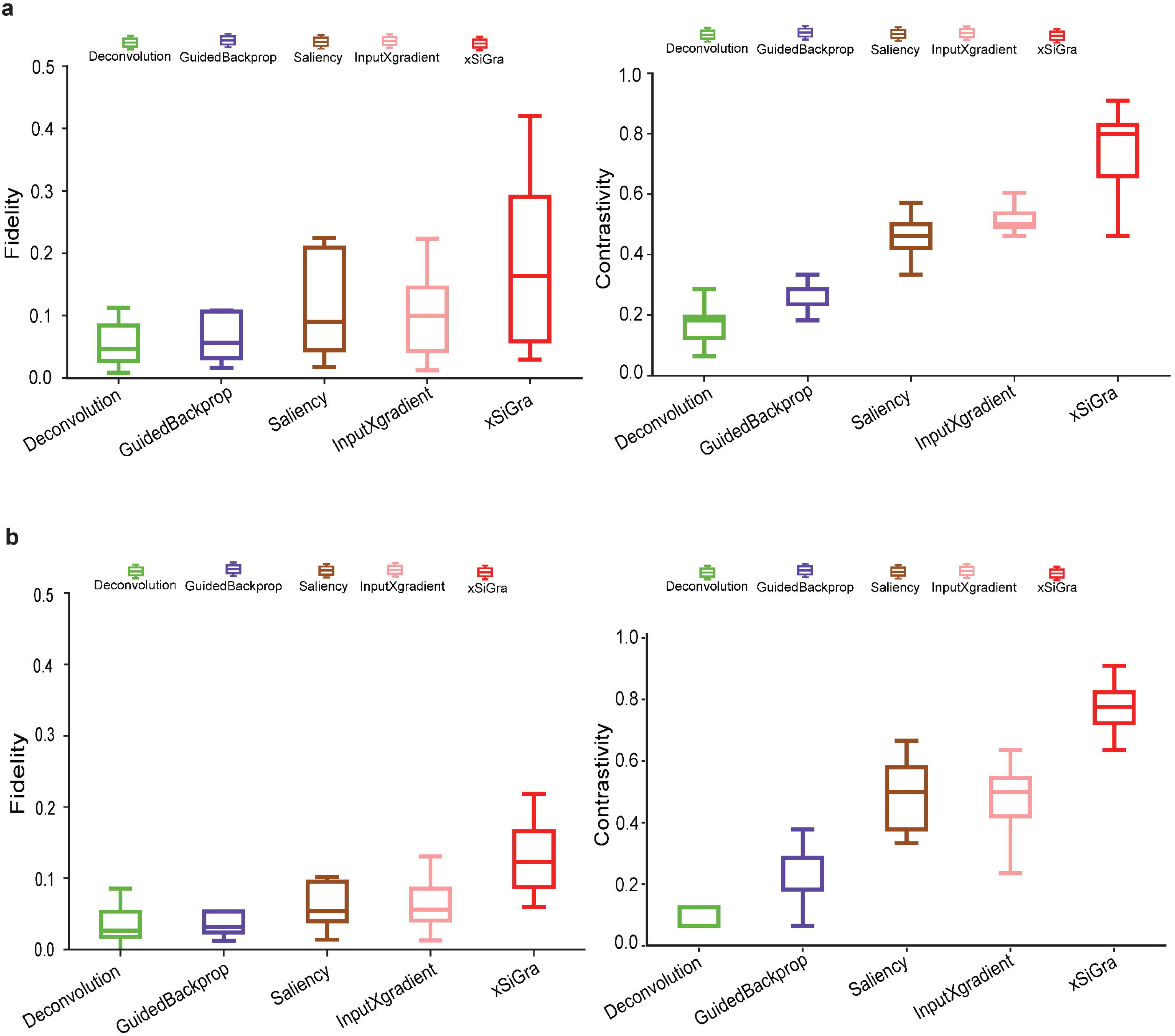
Evaluation results of model explainability. **a** Boxplot of fidelity and contrastivity of Deconvolution, GuidedBackprop, Saliency, InputXGradient, and xSiGra methods on Lung-13 tissue slice. **b** Boxplot of fidelity and contrastivity of those methods on Lung 5-rep3 tissue slice.

### xSiGra identifies important genes and cells for spatial cell types

Given the strong interpretability of xSiGra, it is able to reveal the important genes and cells contributing to the identified spatial cell types. To visualize the cell-level contributions, as shown in **Fig.4a**, we use gradient colors representing cells ranging from low to high contribution for each of the major cell population (tumor, fibroblast, myeloid, and lymphocytes). It is observed that, for a specific cell type, not only the cells within this cell type, but also some neighboring cells contribute to the identification of this specific cell type. To further confirm this observation, we utilize four distinct colors to differentiate cells of the particular cell type that exhibit importance and those not belonging to the specific cell type but still demonstrate importance (**Fig.4b**).

**Fig 4:**
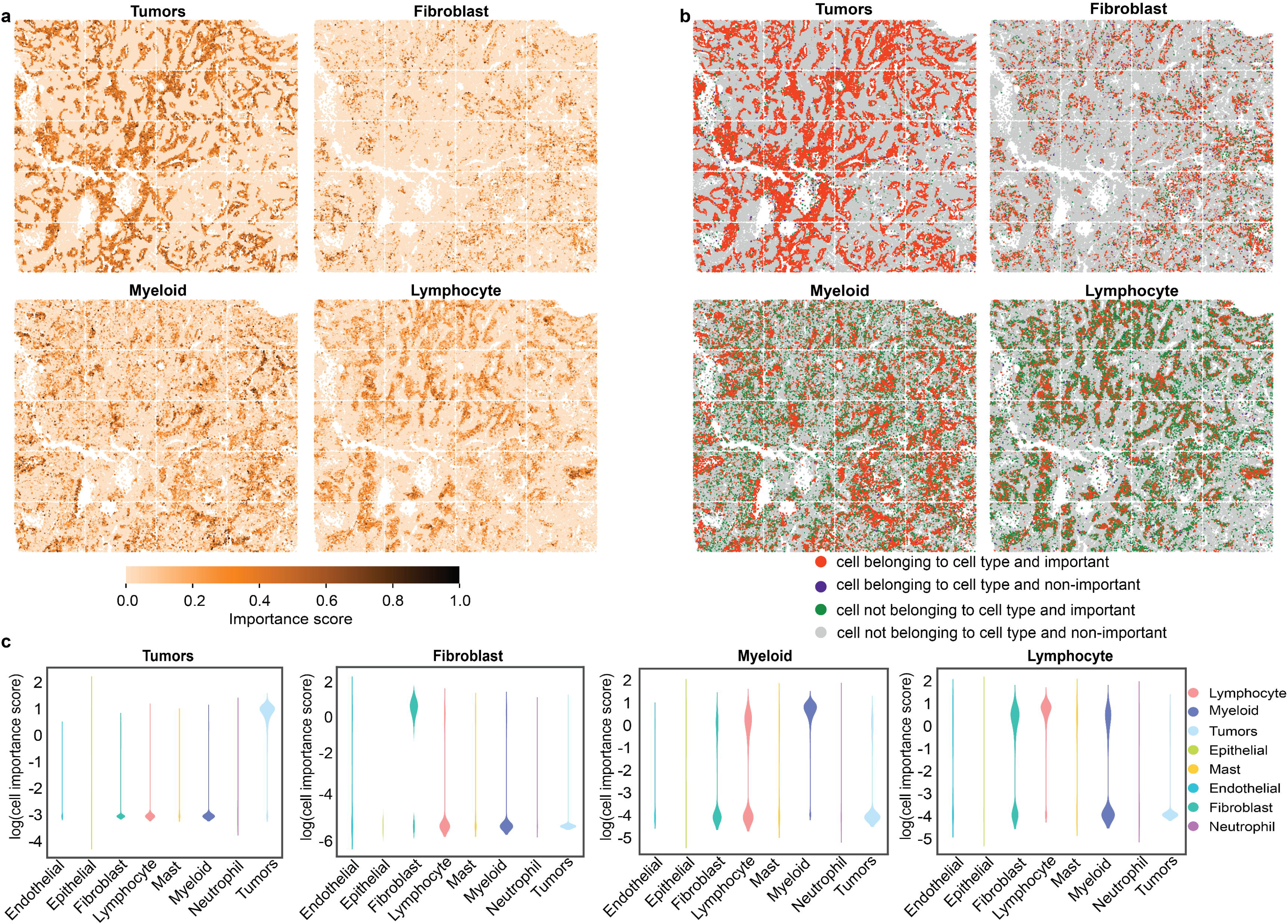
Spatial figure to visualize cell-level contribution in identifying spatial cell types. **a** A color gradient is used to visualize the contribution of cells in identifying spatial cell types. **b** Different color codes are used to distinguish the contribution of cells in identifying the specific cell type. **c** Violin plot to visualize the contribution of cells to each cell type.

Specifically, in the case of tumor cells, the majority of these cells, along with a few non-tumor cells, exhibit significant contributions to the identification of this cell type. However, there are also a few tumor cells that do not contribute to its identification. Similarly, for fibroblasts, most of these cells contribute to their identification along with a few non-fibroblast cells. Comparable patterns are observed for myeloid and lymphoid cells as well. Notably, for each spatial cell type, the cells that contribute to it despite not belonging to that specific cell type are primarily neighboring cells. **Fig.4c** provides quantitative measurements of the contributions of eight different cell types for identifying a particular spatial cell type. In tumor identification, tumor cells make the major contribution, while fibroblasts, along with a few myeloid and lymphocyte cells, contribute significantly to the identification of the fibroblast cell type. Myeloid cells, as well as some lymphocytes and fibroblasts, contribute to the identification of the myeloid cell type. Similarly, lymphocyte cells, along with some myeloid and fibroblast cells, show contributions to the identification of the lymphocyte cell type. Overall, xSiGra offers valuable insights into spatial cell type identification, providing specific information about how cells and their neighboring cells are involved in deciphering spatial cellular heterogeneity.

### Explainable xSiGra uncovers endothelial subset involved in ECM-related interactions

With the interpretable features and their importance scores, we delve deeper into the analysis of cell-cell interactions (CCIs) involving known ligand-receptor (L-R) pairs between adjacent cells (**Materials and Methods**). The CCI analysis reveals significant L-R pairs along with their associated cell types, providing valuable biological insights (**Fig.5a**). **Fig.5b** illustrates the quantification of L-R interactions within CCIs across different cell types, highlighting a strong interaction between endothelial cells and tumor cells, particularly driven by ECM-related interactions. Through interrogation of the ECM-related interactions, notable receptors such as FLT1 and integrins (ITGA2, ITGA3, ITGA6, ITGAV) in endothelial cells were identified. Specifically, in FOV 14, spatial visualization of FLT1 and integrins’ importance scores in endothelial cells is shown (**Fig.5c**), with further visualization across the whole tissue slice and individual FOVs provided in **Supplementary Fig.3**. These observations underscore the diverse importance of FLT1 or integrins among endothelial cells, suggesting their distinct roles in ECM-related interactions. Subsequently, we conduct a comparative analysis between endothelial cells with higher and lower importance scores through differential expressed gene (DEG) analysis, revealing overexpressed genes and enriched pathways in endothelial cells with high FLT1 importance scores (**Fig.5d**), as well as the enriched pathways in those with high integrins importance scores (**Fig.5e**). The top enriched ECM-related pathways emphasizes that endothelial cells with high importance scores play pivotal role in their interaction with tumor cells, indicating a heterogeneous underlying mechanism within this cell type in the complex cellular communications.

**Fig 5:**
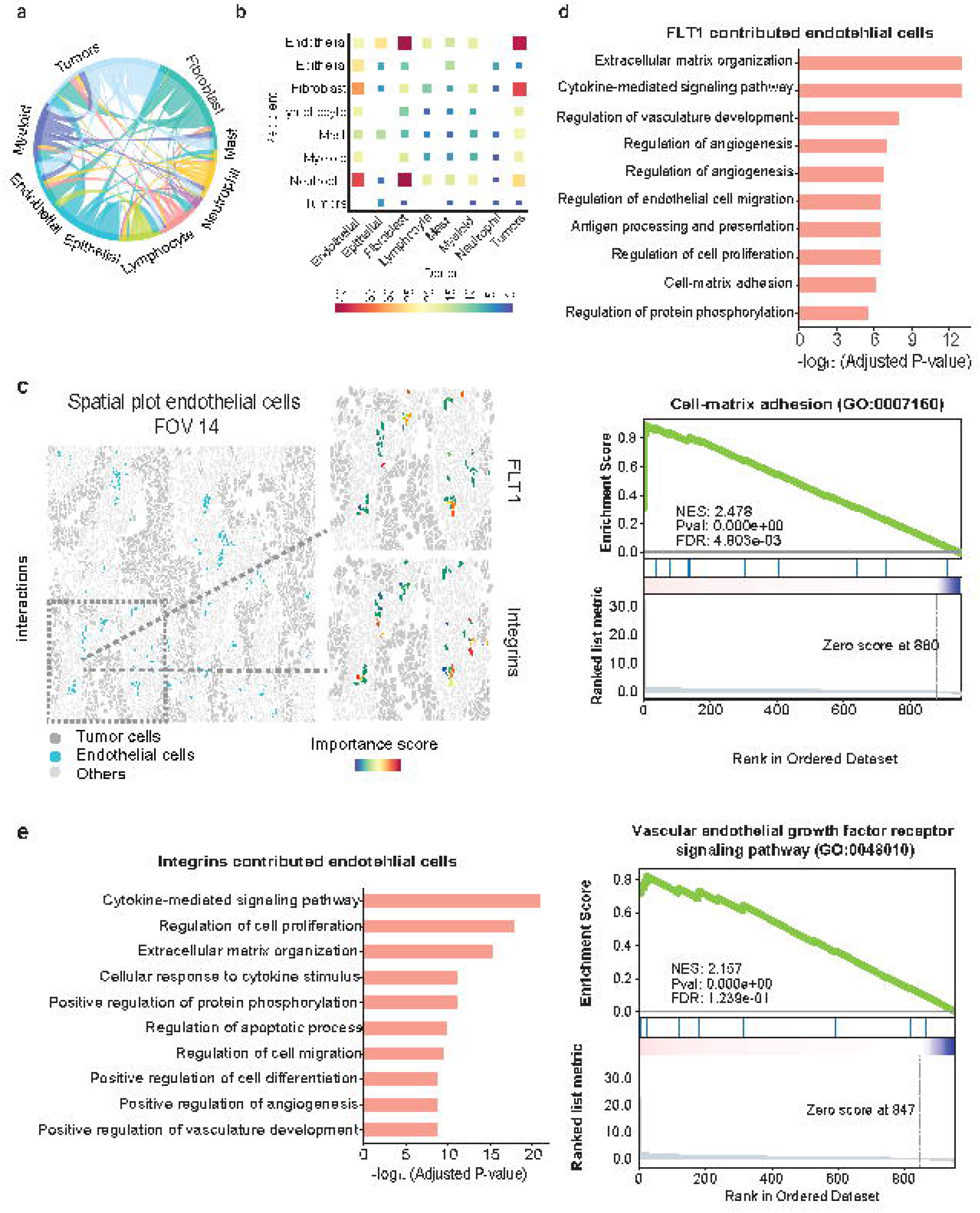
Downstream analysis using the identified important cells and genes. **a** Circular plot of cell-cell interactions. **b** Heatmap of cell-cell interactions with the number of involved L-R interactions from both raw data and importance scores. **c** Spatial visualization of blood vessel endothelial cells in FOV 14. The zoom-in panels show the importance scores of FLT1 and integrins. **d** Gene enrichment analysis of endothelial cells with high importance scores of FLT1. **e** Gene enrichment analysis of endothelial cells with high importance scores of ECM receptors ITGA2, ITGA3, ITGA6, and ITGAV.

## DISCUSSION

Recent advances in spatial transcriptomics technologies have enabled subcellular RNA profiling in tissue slice^1, 28^. Platforms like NanoString CosMx SMI^5^ and Vizgen MERSCOPE^4^ have enabled the spatial profiling of thousands of RNA targets. However, such spatial imaging data faces challenges of missing values and data noise, which can negatively affect downstream analysis such as spatial domain detection^28, 29^. Several deep learning models have been proposed to improve noisy transcriptomics data and perform data analysis^7^, but most of them are black-box approaches that lack transparency and interpretability^18, 30^. To address this challenge, we have proposed xSiGra, which not only accurately identifies spatial cell types and enhances gene expression profiles, but also offers quantitative insights about which cells and genes are important for the identification of spatial cell types, thus making it an interpretable model.

Morphology features have been shown to be linked with gene expression data and combining information from both can lead to better cell prediction performance^31, 32^. xSiGra is engineered to use to the potential of multimodal data such as multichannel cell images and their microenvironment. It includes two major modules, i.e. SiGra+ and graph Grad-CAM. The SiGra+ module is evolved from our recently published SiGra model with an improvement by using a VGG^25^ feature extractor to better leverage the histochemistry images, which are as part of the spatial imaging data, and by introducing a KL divergence loss item to better balance the contributions from transcriptomics and imaging. xSiGra learns the latent space representation, which is used to identify the spatial cell types by Leiden clustering^33^. To make the model explainable, we use a variant of gradient-weighted class activation map (Grad-CAM)^27, 34^. To integrate Grad-CAM with SiGra+, we introduce two linear layers with intermediate non-linear transformation. We train these layers to predict the cell type membership probability through minimizing the cross-entropy loss, using the latent representation as input and vendor-provided cell-types as ground truth. With the new layers replacing Leiden clustering, xSiGra is able to provide the cell and gene importance scores for the spatial cell types. The derived quantitative measures of gene and cell importance for each cell type facilitates downstream analysis like cell-cell interactions and gene enrichment analysis. Importantly, xSiGra is one of the first attempts to quantitatively measure the gene and cell importance to spatial cell types, making it a transparent and trustworthy solution for users.

In addition to the advantages, xSiGra holds considerable promise for continuous improvement in the future. It can be adapted and expanded in several ways. Due to its hybrid architecture, it can easily incorporate emerging modalities in omics data including novel image modalities. It can be further improved by integrating with 3D images and spatial information to use richer information for data analysis. While the current focus of xSiGra is on providing explanations for spatial imaging data, it can be easily extended to provide interpretability for other spatial omics data such as spatial proteomics data. The adaptability of xSiGra for ongoing evolution in spatial technologies is anticipated to empower its utility in the field.

## MATERIALS AND METHODS

### Data processing

Single-cell spatial transcriptomics datasets including 8 lung cancer tissue slices of NanoString CosMx SMI and 12 brain tissue slices of 10x Visium are used. Gene expressions are first normalized by multiplying by 10,000 and then log-transformed. 1) For the NanoString datasets, each whole tissue sample includes a certain number of field of views (FOV). For each cell within a FOV, a 120-pixel x 120-pixel image is cropped with the cell at the center. For NanoString SMI data, a spatial graph is constructed using spatial locations of cells. Cells at a distance < 80 pixels (14.4 μm) are considered as neighbors in the graph. Consider the graph as *G*= (*V,E*) where each *ν* _*i*_ ∈ *V* is a cell node (*i*= 1… *N* and *N* is the total number of cells) and each e_*ij*_ ∈ *E* is the distance between cell ν_i_ and ν_*j*_. Each node has gene expression *g*_*i*_ = {*g*_*i*_,*k*}(*k*= 1 …*K* and *K* are total genes) and image features *m*_*i*_= {*m*_*i*_,*f*}where *f* = 1 … F. 2) In 10x Visium data, for each spot, 50-pixel x 50-pixel image is cropped with the spot as the center. To construct the spatial graph for 10x Visium data, spots at a distance < 150 pixels (16.2 μm) are considered as neighbors in the graph. Similarly, the graph *G*= (*V,E*) has node *ν* _*i*_ ∈ *V* representing a cell (*i*= 1,… *N* and *N* is the total number of cells) and *e*_*ij*_ ∈ *E* representing the edge between the neighboring cell ν_*i*_ and ν_*j*_. Each node has gene expression *g*_*i*_ = {*g*_*i*_,*k*} (*k*= 1… *K* and *K* are total genes) and image features *m*_*i*_= {*m*_*i*_,*f*} where *f*= 1,…,*F*.

### The xSiGra model

The xSiGra model consists of the 1) SiGra+ to reconstruct gene expressions, and 2) graph gradient-weighted class activation mapping (graph Grad-CAM) model to identify interpretable features.

1) SiGra+

SiGra+ is a hybrid graph encoder-decoder framework, consisting of three modules: i) the image module, ii) the gene module, and iii) the hybrid module. Specifically, i) in the image module, for each node ν_i_ in the graph, features from the images *m*_i_ are first extracted using a pre-trained VGG-16 network. The extracted image features are then passed through graph transformer layers^35, 36^ to obtain the latent space Z_*m,i*_,. This latent space representation further goes through additional graph transformer layers to reconstruct the gene expressions of node ν_i_. Here 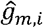 represents the reconstructed gene expression from Z_*m,i*_. ii) In the gene module, for each node ν_i_ in the graph, gene expression features are passed through graph transformer layers to obtain the latent space Z_*g,i*_. This latent space representation goes through graph transformer layers *T* ^(*l*)^(*l* denotes the *l*’th layer) to reconstruct the gene expression counts for the node ν_i_. Here 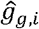 represents the reconstructed gene expression from Z_*g,i*_ iii) In the hybrid module, the two latent space features *Z*_*m,i*_, *Z*_g,i_ are concatenated and provided as input to graph transformer layer which projects it to new latent space *Z*_*h,i*_. The hybrid embedding *Z*_*h,i*_ is further passed through graph transformer layers to reconstruct the gene expressions of the node ν_*r*_. Here 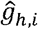 represents the reconstructed gene expression from hybrid embeddings.

With the input graph *G*= (*V,E*), the multimodal features (image and gene expression) of neighboring nodes also contribute to the reconstructed gene expressions of node ν_*i*_.

The hybrid graph encoder-decoder is trained to learn gene expressions with the loss function (*L*) as:

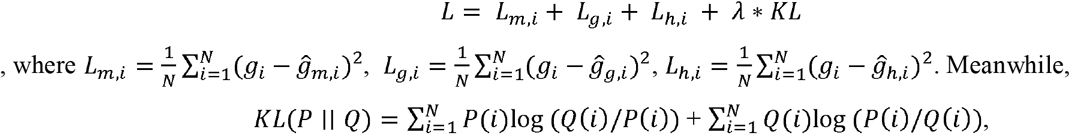

where *P* and *Q* are reconstructed gene expressions pairs 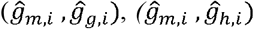 and 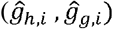. Here, λ is chosen as 0.001 using hyperparameter tuning.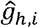 is used as the final reconstructed gene expression and *z*_*h,i*_ is used for clustering to identify spatial cell types using the Leiden algorithm ^33^.

2) Graph Grad-CAM model

xSiGra uses a novel post hoc graph gradient-weighted class activation mapping (graph Grad-CAM) model to reveal the importance of genes and cells for biological functions of interest, such as the identified spatial cell types. Since the spatial cell types are identified through unsupervised clustering of the latent space *z*_*h,i*_, it is challenging to directly interpret the results. Herein, we first train an auxiliary AI classifier component to map spatial cell types to the corresponding ground truth, then use graph Grad-CAM to explore the importance of genes and cells for the predicted probabilities of spatial cell types.

For the auxiliary classifier component, we choose a dense multilayer perceptron as the classifier, with intermediate non-linear ReLU^37^ activations as the hidden layers, and a log softmax activation layer to predict probabilities of spatial cell types. The log softmax activation is used as it shows better stability and faster gradient optimization over softmax activation^38^. Briefly,

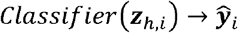

where, for cell *i*, 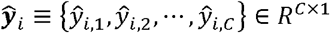is a vector of the predicted probabilities of cell *i* belonging to each of the *c* spatial cell types. For the NanoString CosMx data, the vendor-provided cell type annotations *y*_*i*_ ≡ {*y*_*i*,1_, *y*_*i*,2_,·, *y*_*i,C*_} are used as ground truth for training the classifier, where *y*_*i,c*_ ∈{0,1} is the cell type label and for each cell there is only one non-zero label. The cross-entropy loss used for training is:

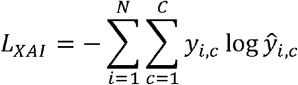

This classifier maps each cell from the latent space *z*_*h,i*_ to its corresponding cell type and thus allows interpreting the gene and cell importance.

Then we developed a novel graph Grad-CAM algorithm to explore the contributions of each gene in each cell to the prediction of each cell type with backpropagation. Consider the spatial graph with *N* cells (nodes) and *K* genes, the *k*-th feature at *1*-th layer in the SiGra+ module for cell *i*, is denoted as 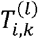. For spatial cell type *c*, its specific weights for gene *k* (or latent feature *k* for *1*> 0) of cell *i*, at the *1*-th layer *T*^(*l*)^ of the SiGra+ gene module is given by:

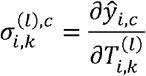

The feature importance at the *1*-th layer is determined through backward propagation:

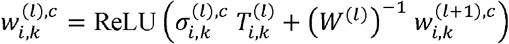

where *W*^(*l*)^ is the learned parameter of *1*-th transformer layer *T*^(*l*)^. The importance score of gene k in cell, for spatial cell type *c* is determined as:

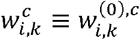

Similarly, the importance score of cell *i*, for spatial cell type *c* is determined as:

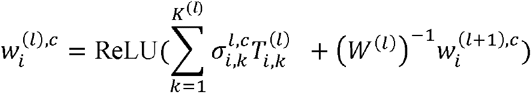

where *K*^(*l*)^ is the number of latent features at the *1*-th layer, and *K*^(0)^ = *K*. For simplicity, in xSiGra, we only use the input layer, i.e., *1*= 0, to compute gene and cell importance. Specifically, the importance of gene *k* in cell *i*, is computed as:

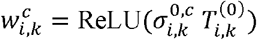

and the importance of cell *i*, for determining cell type *c* is:

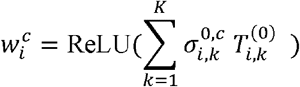

where 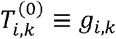 is gene expression count at layer 0 for cell *i*, and gene *k*. Thus, the importance of genes and cells for each spatial cell types are determined. In this work, through fine-tuning, we determine the hyperparameters as: 2 layers for the auxiliary classifier with the dimensions of each layer as: 1024 and 8.

## Performance benchmarking

For benchmarking on the accuracy of identified spatial cell types, we compare our method with six existing methods, including SiGra^9^, SpaGCN^10^, stLearn^11^, BayesSpace^12^, STAGATE^13^, Seurat^14^ and Scanpy^15^. The performance of different methods is evaluated using the adjusted rand index (ARI) score. That is, suppose 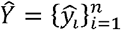 represent the spatial cell types and 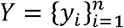 represent the ground truth, i.e., *k* cell types or spatial domains from n cells or spots, ARI is calculated as:

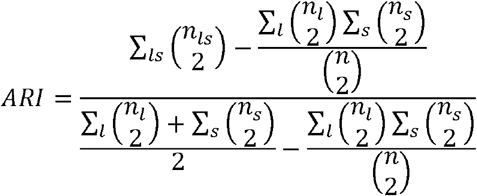

where *1* and *s*, denote the *k* cell types, 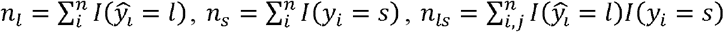 and *I*(*x* = *y*)=1 when *x*= *y*, else *I*(*x* = *y*) = 0

To evaluate the explainable power of xSiGra, we have compared it with the existing explainable methods, namely Saliency^20^, InputXGradient^22^, GuidedBackprop^21^, and Deconvolution^23^. The model explainability is measured with two metrics, i.e., fidelity score and contrastivity score. Specifically, 1) fidelity score is computed as the differences between the model’s C-index score (Area Under the Receiver Operating Characteristic Curve, AUROC) when all genes are used and when the top 30 important genes are masked. The fidelity score is computed separately for different spatial cell types and the median value of fidelity scores is used for comparison. A higher fidelity score indicates better explainable capability. 2) Contrastivity score is computed as 1-Jaccard similarity for each pair of different spatial cell types. Specifically, if *I*_*i*_ represents the list of top 30 important genes identified for cell type *i*,, *I*_*j*_ is the list of top 30 important genes identified for cell type *j*, then the contrastivity score for cell types *i*, and *j* is computed as: 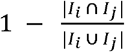. The median value of contrastivity scores across all pairs of spatial cell types is used for performance evaluation.

### Enrichment analysis

Gene sets of the Reactome and KEGG pathway database are downloaded from the MSigDB Collections^39, 40^. Functional enrichment based on the above respective databases is assessed by hypergeometric test, which is used to identify a priori-defined gene set that shows statistically significant enrichment. The test is performed by the Scanpy package. We further correct the P-values by Benjamini-Hochberg and those with less than 0.05 are considered as statistically significant.

### Cell-cell interaction analysis

To conduct the cell-cell interaction analysis we first built a spatial graph using cell location. Cells at a distance < 80 pixels (14.4 μm) are considered as neighbors in the graph. Then, we use a known set of ligand-receptor pairs^41^ to compute interaction scores for all cells in the spatial graph. For each neighbor cell pair *ν*_*i*_ and *ν*_j_ we have *L*_*i*_ and *L*_*j*_ as the ligand-related expression or importance score in cell *ν*_*i*_ and cell *ν*_*j*_, while *R*_*i*_ and *R*_*j*_ as the receptor-related expression or importance score in cell *ν*_i_ and cell *ν*_j_ respectively. The interaction scores are computed as score1 = *L*_*i*_ × *R*_*j*_ and score2 = *L*_*j*_ × *R*_*i*_. Next, the interaction scores for each unique ligand gene-cell type and receptor gene-cell type neighbor pairs are aggregated using average. To select the statistically significant interactions, we compute z-scores and choose the interactions having FDR adjusted p-value below 0.05. The significant interactions identified using the analysis are further used to gain biological insights.

---

**Supplementary Fig 1:**
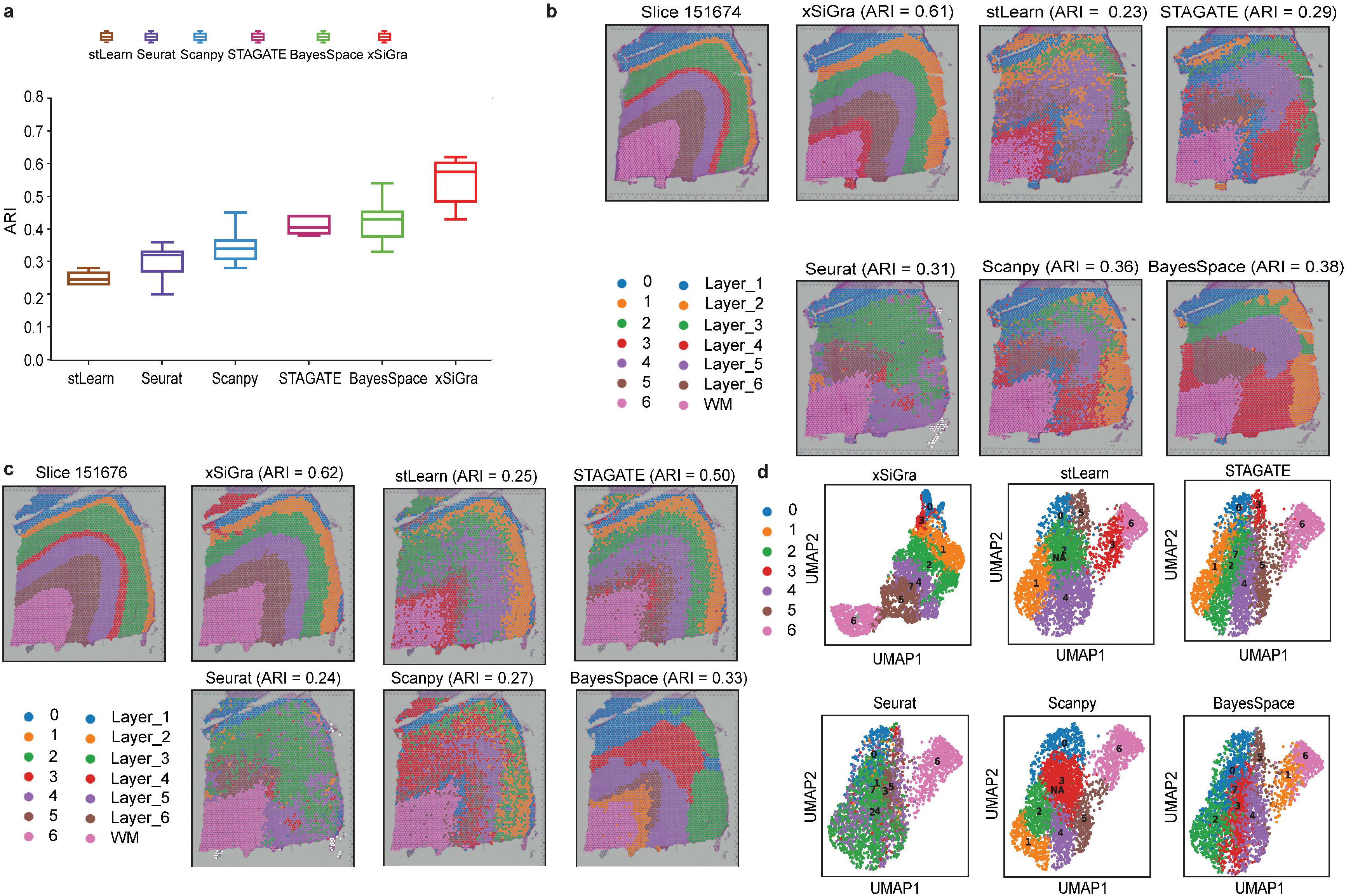
ARI results of benchmarking methods on 10x Visium slices. **A** Boxplot of adjusted rand index for 12 10x Visium datasets is shown for the methods, i.e. stLearn, STAGATE, Seurat, Scanpy, BayesSpace and our method. **b** Spatial domains of ground truth and those identified by each method are shown for slice 151507. **c** Spatial domains of ground truth and those identified by each method is shown for the slice 151676. **d** UMAP visualizations of latent embeddings by stLearn, Seurat, Scanpy, and our method are shown for slice 151576.

**Supplementary Fig 2:**
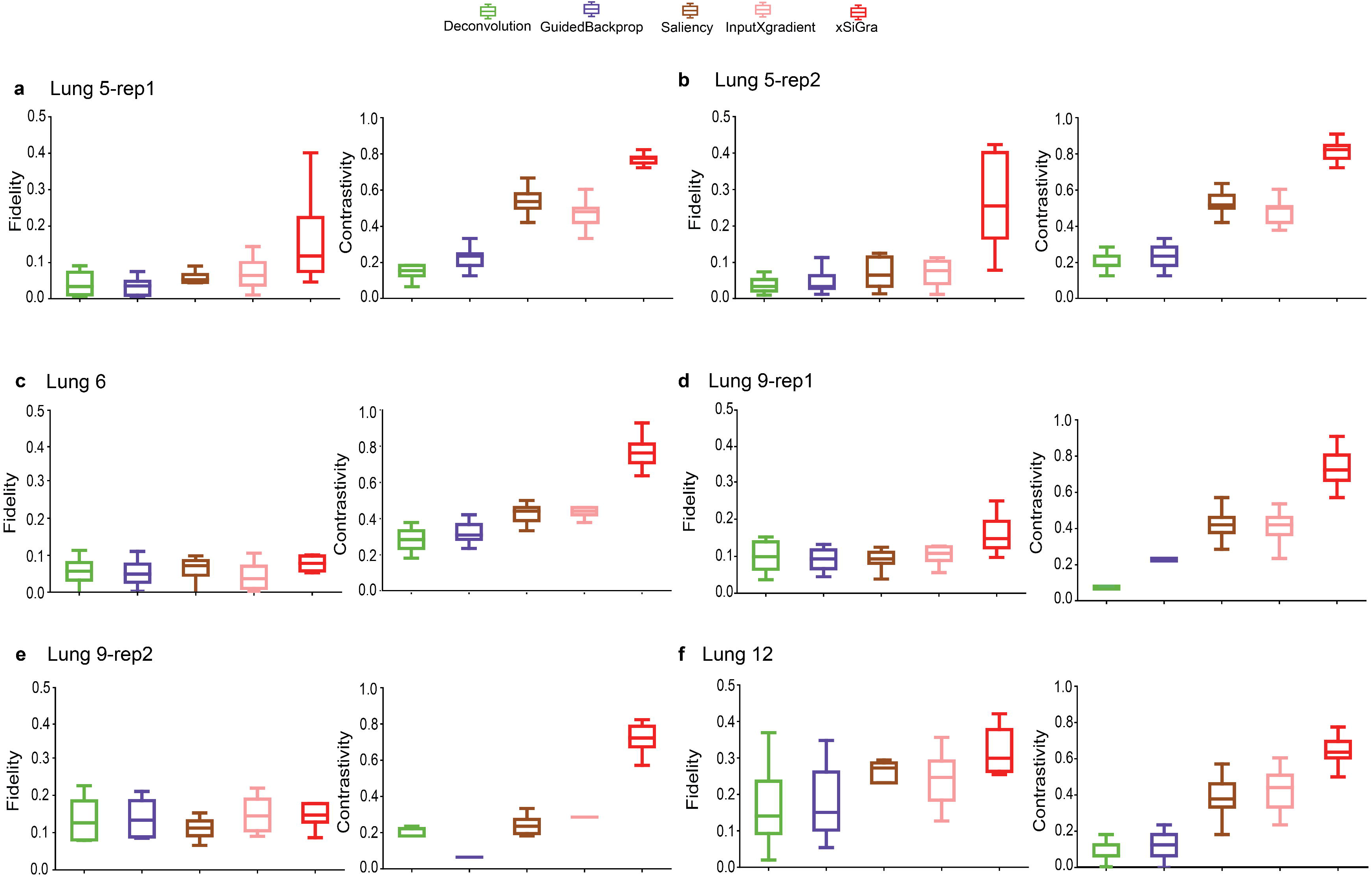
Evaluation results for model explainability on different lung cancer slices. Boxplot of fidelity and contrastivity for Deconvolution, GuidedBackprop, Saliency, InputXGradient, and xSiGra algorithms for Lung 5-rep1, Lung 5-rep2, Lung 6, Lung 9-rep1, Lung 9-rep2, and Lung 12 tissue samples.

**Supplementary Fig 3:**
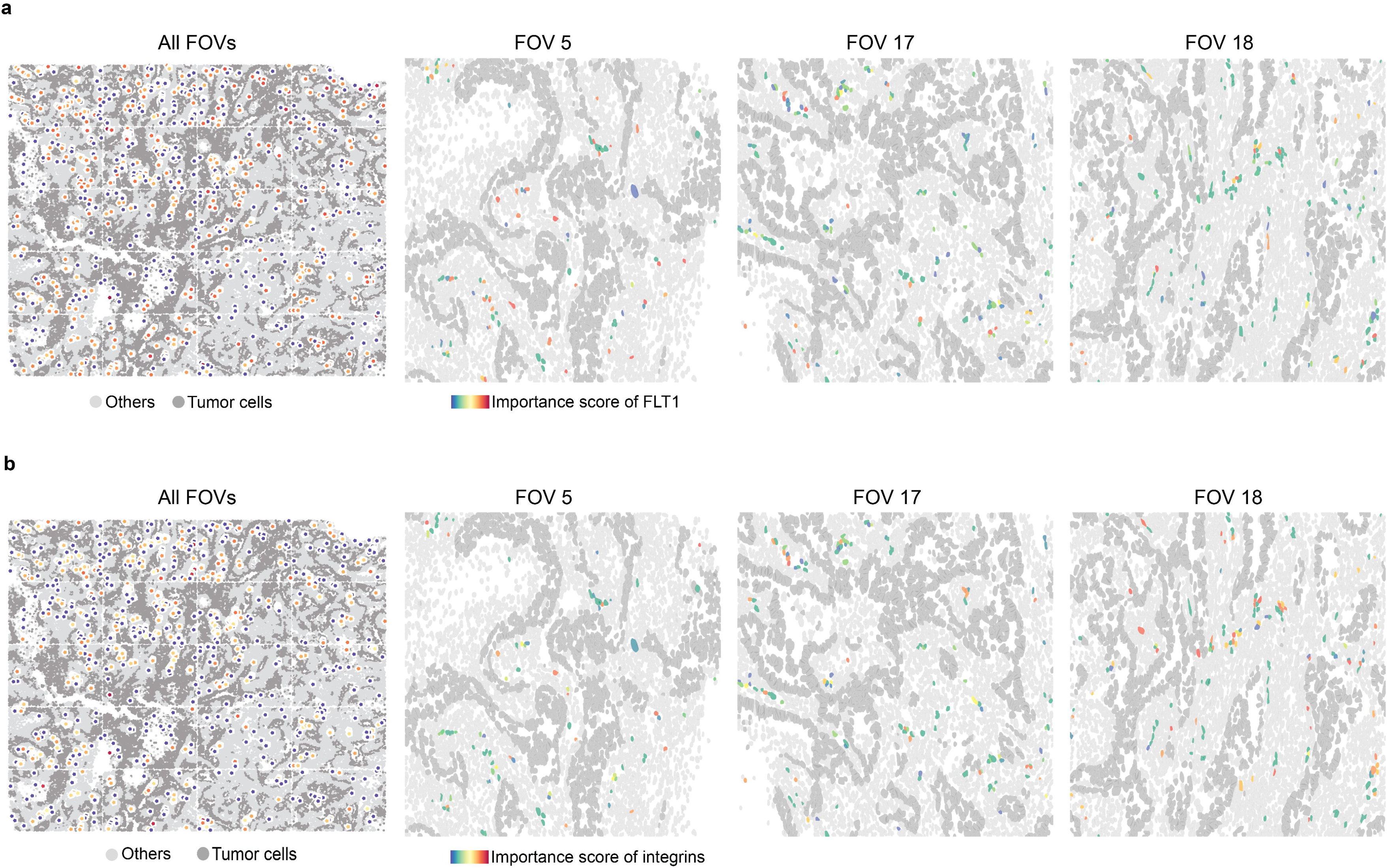
Spatial visualization of endothelial cells. Spatial visualization of blood vessel endothelial cells in all FOVs. The right panels show the importance scores of FLT1 (**a**) and integrins (**b**).

## DATA AVAILABILITY

In this study, we have used the NanoString CosMx SMI non-small-cell lung□cancer (NSCLC) formalin-fixed paraffin-embedded (FFPE) Dataset and 10x Visium datasets from human dorsolateral prefrontal cortex (DLPFC)^42^. NanoString dataset contains 8 different samples from 5 NSCLC tissues, Lung 5-rep1, Lung 5-rep2, Lung 5-rep3, Lung6, Lung 9-rep1, Lung 9-rep2, Lung-12 and Lung-13. For each dataset gene expression data and multichannel images are provided with 5 channels: MembraneStrain, PanCK, CD45, CD3, and DAPI. For each staining, a rich grey-scale image composed of a combination of multiple FOVs is also provided. Along with that metadata about cells such as identified cell coordinates, cell area, width, height is provided. Lung-13 sample consists of 20 field of views (FOVs) and 77643 cells. The cells were grouped into 8 cell types: tumor, fibroblasts, lymphocyte, mast, neutrophil, endothelial and epithelial cells. 10x Visium dataset consists of 12 samples, 151676, 151675, 151674, 151673, 151672, 151671, 151670, 151669, 151510, 151509, 151508, 151507, with up to 6 cortical layers and white matter manual annotation. The hematoxylin and eosin (H&E) images of the tissue sections along with the transcriptomics data are provided.

## CODE AVAILABILITY

The xSiGra model is provided as an open-source python package in GitHub (https://github.com/QSong-github/xSiGra), with detailed manual and tutorials.

